# A shared circuit architecture for multiscale signal recognition in crickets

**DOI:** 10.64898/2026.09.28.754905

**Authors:** Winston Mann, Daesung Cho, David A Gray, R Matthias Hennig, Jan Clemens

## Abstract

Communication signals often contain behaviorally relevant structure on multiple temporal scales. Short-timescale features are commonly thought to be recognized first and then integrated into longer-timescale representations. Whether a shared circuit can instead recognize multiple timescales remains unclear. Cricket calling songs provide a tractable test because millisecond-scale pulses are grouped into chirps lasting hundreds of milliseconds. We combined female phonotaxis with a mechanistic model of the identified pulse-recognition network to ask whether this circuitry can also generate chirp selectivity. In *Gryllus bimaculatus*, the model, originally parameterized from pulse-scale electrophysiology and behavior, recovered the main boundaries of chirp preference without fitting to chirp behavior. Coincidence detection generated the minimum chirp-duration requirement, whereas suppressive computations in the network’s input neuron shaped the upper boundaries for chirp duty cycle and period. Across 14 species, varying these suppressive computations captured major differences in chirp tuning. These findings reveal a temporally interleaved serial circuit in which long-timescale selectivity begins upstream of completed short-timescale selectivity, offering an alternative to both conventional short-first hierarchies and parallel pathways.

## Introduction

Communication signals are hierarchically organized, with behaviorally relevant information distributed across multiple temporal scales. Human speech consists of phonemes, syllables, words and sentences, birdsong is organized into syllables and motifs, and many frogs and insects produce chirps composed of stereotyped sound pulses (Garcia-Lazaro et al., 2013; Gerhardt and Huber, 2002) (Fig. 1A, B). Successful communication therefore requires nervous systems to recognize temporal patterns spanning tens of milliseconds to several seconds. While substantial progress has been made in understanding how temporal features are extracted on individual timescales (Creutzig et al., 2010; Machens et al., 2001; Schöneich et al., 2015; Lewicki and Konishi, 1995; Moore and Woolley, 2019; García-Rosales et al., 2018), much less is known about how recognition across multiple timescales is organized within neural circuits and how these computations evolve as communication signals diversify (Rose et al., 2015; Clemens et al., 2021).

**Figure 1.**
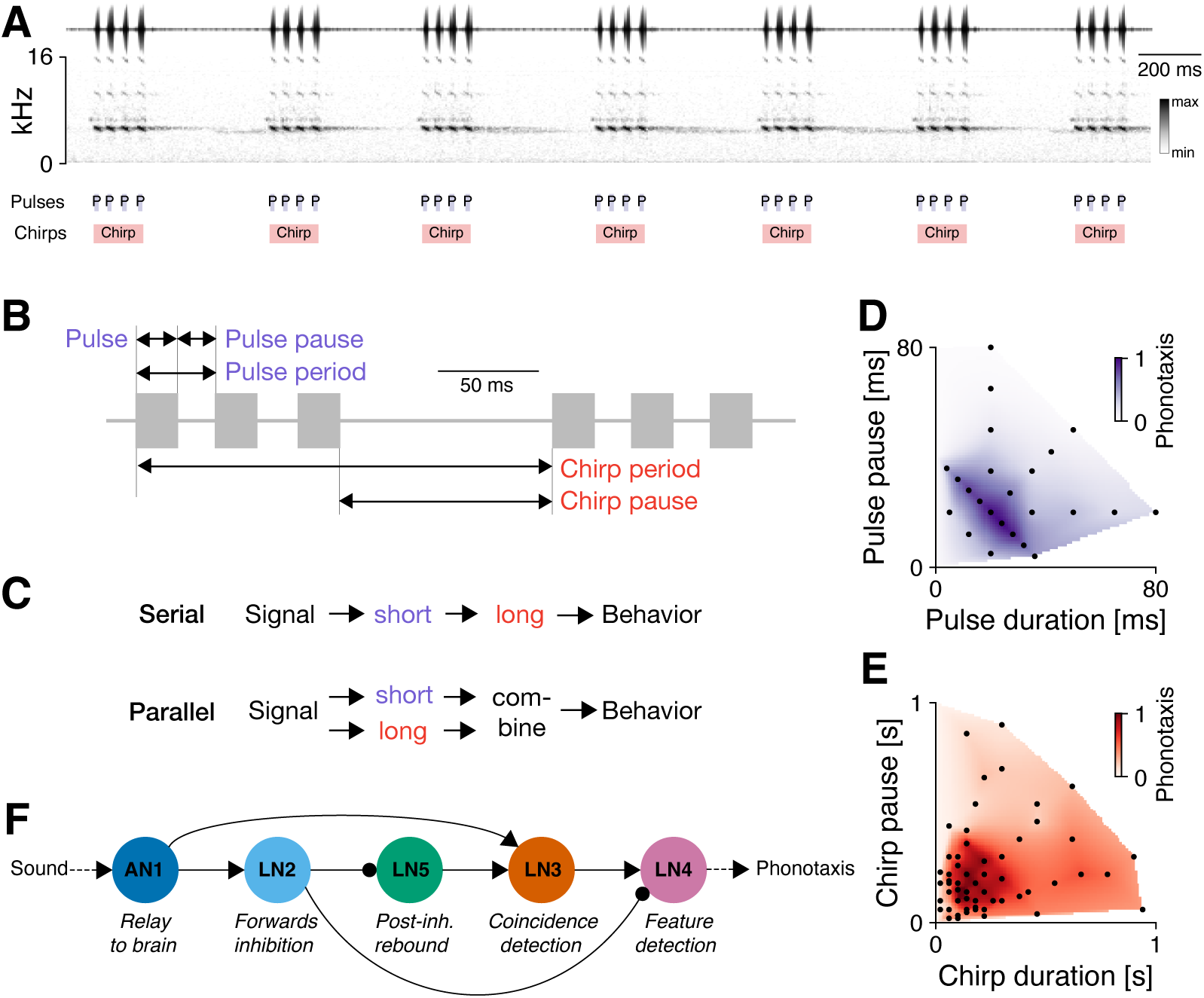
Multiscale calling-song structure and recognition in crickets. **A** Hierarchical temporal organization of cricket calling song, from short constituent elements (pulses (P), purple) to longer units (chirps, red). The top shows the waveform and the bottom the spectrogram of calling song from *G. bimaculatus* (see color bar). **B** Structure of calling song at the pulse and chirp scales in *G. bimaculatus*. Pulses have stereotyped durations (20 ms) and are separated by stereotyped pulse pauses (20 ms). Several pulses are produced in short bursts called chirps with a specific chirp duration (80–160 ms) and chirp pause (120–270 ms). Both timescales can also be characterized via the derived measures *period* (duration plus pause) and *duty cycle* (duration divided by period). See Table S1 for song statistics. **C** Candidate architectures for multiscale feature recognition: serial processing (top) in which short (purple) timescale features are integrated into long (red) timescale features and parallel pathways (bottom) in which short and long timescales are processed independently and subsequently combined. **D, E** Female preferences for features at each timescale can be measured using phonotaxis experiments. *G. bimaculatus* pulse (D) and chirp (E) preference fields reveal well-defined regions in pulse and chirp space preferred by females. Points mark tested stimuli and color codes interpolated preference values. **F** Pulse-recognition network in *G. bimaculatus*. AN1 provides direct excitation to LN2 and LN3. LN2 inhibits LN5 and LN4. LN3 only fires strongly when a post-inhibitory rebound from LN5 coincides with AN1 input. LN3 excites the feature detector LN4, whose output predicts the pulse preferences in (D).

A central question is whether recognition across temporal scales is organized serially or in parallel (Fig. 1C). A widely held view is that progressively longer temporal features are computed hierarchically from shorter ones. In speech perception, auditory representations are commonly described as progressing from acoustic features to phonemes, syllables, words and sentences (Friederici, 2012; Giraud and Poeppel, 2012; Ding et al., 2016). Likewise, theoretical, neurophysiological and imaging studies describe progressively longer temporal receptive windows and intrinsic timescales from early sensory areas to higher-order association cortex (Kiebel et al., 2008; Hasson et al., 2008; Lerner et al., 2011; Honey et al., 2012; Murray et al., 2014; Chaudhuri et al., 2015; Hasson et al., 2015; Runyan et al., 2017). An alternative view is that different temporal features are extracted by parallel pathways operating over distinct integration windows before converging onto downstream decision circuits.

Variants of both architectures have been proposed in compact auditory systems. In frogs, interval-counting neurons integrate a threshold number of correctly timed pulses over tens to hundreds of milliseconds, consistent with a long-term integration stage operating on shorter pulse relationships (Alder and Rose, 1998; Edwards et al., 2002; Edwards et al., 2007; Rose et al., 2015). In grasshoppers, song recognition has instead been explained by parallel detectors of pulse duration, pause duration and pulse number whose outputs converge onto a common readout (Creutzig et al., 2009; Creutzig et al., 2010; Clemens and Ronacher, 2013; Ronacher et al., 2014). Thus, both serial accumulation and parallel feature extraction are biologically plausible solutions. Direct evidence distinguishing how these principles are organized across a circuit of identified neurons nevertheless remains limited (Giroud et al., 2024; Chen et al., 2024).

The cricket auditory system provides a tractable model for addressing this question. Male calling songs consist of brief pulses grouped into longer chirps (Fig. 1A, B), and female phonotaxis depends on temporal structure at both scales (Fig. 1D, E) (Kostarakos et al., 2009; Grobe et al., 2012; Hennig et al., 2016). Previous behavioral work showed that chirp duty cycle and chirp period are the principal long-timescale cues and that unattractive structure at either scale can suppress responses to otherwise attractive patterns. Strong phonotaxis therefore generally requires suitable pulse and chirp structure, consistent with a weighted AND-like interaction between the two timescales (Grobe et al., 2012; Meckenhäuser et al., 2013). This interaction, however, is compatible with both serial processing and parallel pathways followed by convergence, and therefore does not reveal where and how the two timescales are computed. Because pulse and chirp parameters have diversified extensively among closely related species (Grobe et al., 2012; Blankers et al., 2015; Hennig et al., 2016; Gray et al., 2016), cricket song recognition has become a classic model for investigating both the neural basis and the evolution of acoustic communication (Pollack, 2000; Gerhardt and Huber, 2002; Kostarakos and Hedwig, 2014; Schöneich, 2020; Mann et al., 2025).

The neural mechanisms underlying pulse recognition have been characterized in detail (Kostarakos and Hedwig, 2012; Schöneich et al., 2015). Auditory information is relayed through a small network of identified interneurons that implements delay lines, post-inhibitory rebound, coincidence detection and suppressive computations to generate selectivity for pulse timing (Fig. 1F). Computational models based on this circuit accurately reproduce female pulse preferences and explain the diversity of pulse tuning observed across cricket species through quantitative changes in this recognition network (Clemens et al., 2021; Mann et al., 2025). In contrast, the neural basis of chirp recognition remains poorly understood. Because chirps extend over hundreds of milliseconds, they are generally assumed to require additional computations downstream of the pulse recognition network (Grobe et al., 2012; Gray, 2022).

Here we combine behavior with an independently constrained circuit model to test the alternative hypothesis that chirp selectivity can emerge from computations present within the pulse-recognition network. We first characterize behavioral responses of *G. bimaculatus* females across the space of chirp duration, period and duty cycle and show that these responses can be summarized by a small number of response boundaries in song feature space. We then test whether an established computational model of the pulse-recognition network reproduces these boundaries without fitting to chirp stimuli. Systematic computational perturbations indicate that, in the model, chirp selectivity arises predominantly from suppressive computations in the peripheral, ascending auditory neuron AN1, whereas pulse selectivity continues to emerge through downstream coincidence-detection circuitry. Thus, the model implements an organization distinct from both the prevailing short-first, long-later serial model and a parallel architecture: the principal computations shaping recognition on the longer timescale are located before those responsible for recognition on the shorter timescale. Finally, we show that quantitative variation in these early suppressive computations reproduces major differences in chirp tuning across cricket species. Together, behavior and the independently constrained circuit model support a temporally interleaved organization of multiscale recognition within a compact auditory network. Our analysis tests the computational sufficiency of this organization and generates predictions for its direct physiological evaluation.

## Results

### Female chirp preferences in *G. bimaculatus* are defined by three simple behavioral boundaries

To characterize how *G. bimaculatus* females evaluate chirp structure, we measured phonotactic responses to synthetic songs that systematically varied in chirp duration and pause while keeping pulse parameters fixed at their preferred values (Grobe et al., 2012). Female responses formed a continuous preference field, with strong phonotaxis restricted to a well-defined region of the chirp parameter space (Fig. 2A). We next asked whether this two-dimensional response field could be summarized by a small number of simple behavioral rules. To this end, we examined one-dimensional transects through the preference field corresponding to the principal chirp parameters (Fig. 2B–F, S1). Chirp duration exhibited a sharp lower threshold at 40 ms (Fig. 2B). Given the fixed pulse duration (20 ms) and pause (20 ms), this threshold corresponds to the transition from one-pulse to two-pulse chirps (from 20 ms to 60 ms) (Fig. 2C). In contrast, responses to chirp period were band-pass (Fig. 2E). Female phonotaxis was restricted to periods between approximately 150 and 600 ms, rejecting both very short chirps with short chirp pauses and very long chirps with long pauses. Finally, we quantified responses as a function of chirp duty cycle, defined as chirp duration divided by chirp period (Fig. 2F). Again, tuning was band-pass, with female phonotaxis restricted to duty cycles between approximately 0.1 and 0.8, rejecting both very sparse and very dense songs. The lower boundary corresponds to sparse chirps with fewer than two pulses (Fig. 2B, C). The upper boundary implies that females reject songs in which long chirps are separated by relatively brief pauses. Together, these analyses show that the two-dimensional chirp preference field in *G. bimaculatus* can be reduced to three major behavioral boundaries: a minimum chirp duration/period, a maximum chirp period, and a maximum duty cycle.

**Figure 2.**
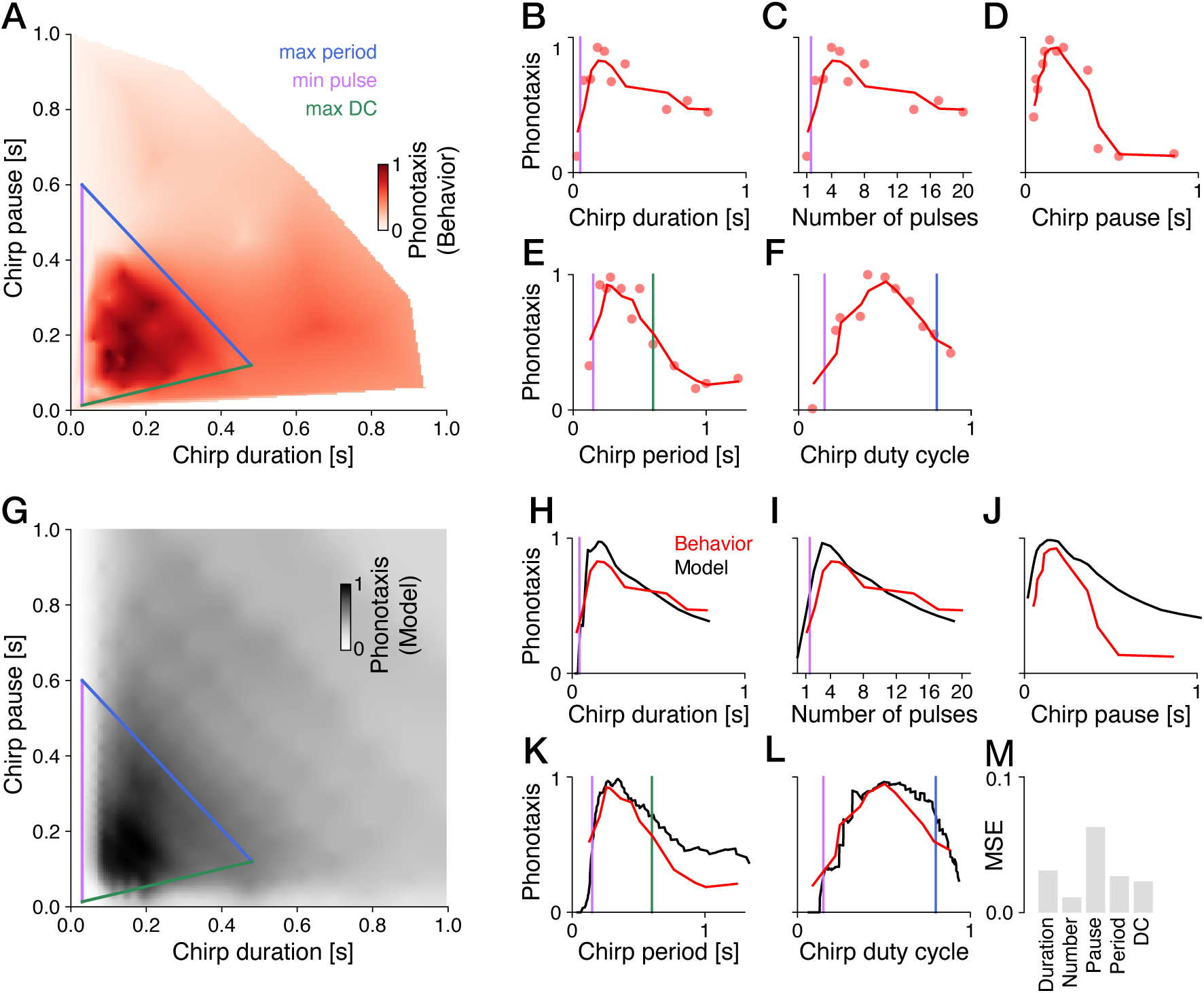
Three boundaries describe behavioral selectivity for chirp parameters in *G. bimaculatus* and are reproduced by a pulse-recognition model. A Phonotaxis across chirp duration and pause. Lines mark the minimum chirp-duration/pulse-number boundary (approximately 40 ms, between one-and two-pulse chirps, purple), maximum chirp-period boundary (600 ms, blue), and maximum chirp-duty-cycle boundary (0.8, green). Phonotaxis was normalized to a maximum of 1. **B–F** Phonotaxis as a function of chirp duration (B), pulse number (C), chirp pause (D), chirp period (E) and chirp duty cycle (F). Red points and curves show measured responses and smoothed trends, respectively, and vertical lines mark the corresponding boundaries. Responses were normalized to their maxima over the full stimulus set. Transects defined in Fig. S1. **G** Normalized model response field with the same boundaries overlaid. The average firing rate of LN4 in the model was taken to be proportional to the modeled phonotaxis response. **H–L** Behavioral (red) and model (black) tuning for chirp duration (H), pulse number (I), chirp pause (J), chirp period (K), and chirp duty cycle (L). Behavioral and model responses were normalized to their respective maxima. **M** Mean-squared error (MSE) between behavioral and model tuning curves.

### Computations in the pulse recognition network explain chirp preferences

The three behavioral boundaries provide a compact description of chirp selectivity and a benchmark for mechanistic models. We therefore asked whether the established computational model of the cricket pulse recognition network, whose parameters were originally determined from pulse-scale electrophysiology (Clemens et al., 2021), could also account for chirp preferences without fitting to chirp stimuli.

We simulated responses of the published model across the chirp-duration and chirp-pause ranges sampled in the behavioral experiments (Fig. 2G). The model was developed to reproduce electrophysiological responses to single chirps and exhibited unstable response dynamics when driven by the much longer stimulus sequences considered here. We localized this effect to divisive normalization in AN1, which reduces a response by dividing it by a gain-control signal that reflects recent activity. This signal accumulates during repeated pulses, so later pulses evoke smaller responses. In the original model, divisive normalization was too strong. We therefore rectified the AN1 signal before divisive normalization and reduced the strength of AN1 divisive normalization. These two implementation changes stabilized responses to long stimulus sequences while leaving pulse tuning and the model’s qualitative chirp selectivity largely unchanged (see Methods, Fig. S2, S3).

This model reproduced the overall structure of the behavioral preference field (Fig. 2G–M), including all three behavioral boundaries identified above. Model responses exhibited a sharp lower threshold at two pulses per chirp (Fig. 2H, I), band-pass tuning for chirp period with both lower and upper boundaries (Fig. 2K), and declining responses at high chirp duty cycles (Fig. 2L). Model and behavior agreed well (Fig. 2M). The largest discrepancy occurred for long chirp pauses, for which the model predicted stronger phonotaxis than observed experimentally (Fig. 2J). Nevertheless, the locations of the behavioral boundaries and the overall shapes of the tuning curves were accurately reproduced.

Together, these results show that computations previously identified as sufficient for pulse recognition can also explain the principal features of chirp selectivity. Neither a dedicated chirp-recognition stage nor fitting to chirp behavior was required. This is notable because chirps extend over hundreds of milliseconds, whereas the model parameters were originally constrained by responses on the pulse timescale of tens of milliseconds. We therefore next asked which computations within the pulse recognition network generate each of the three behavioral boundaries.

### Coincidence detection generates the minimum chirp duration threshold

The behavioral threshold of two pulses per chirp suggests that the minimum chirp duration arises from coincidence detection, which requires responses to successive pulses to coincide and therefore at least two pulses per chirp (Schöneich et al., 2015). To test this idea, we compared model responses to one-and two-pulse chirps across successive stages of the network.

In neurons upstream of coincidence detection—AN1, LN2, and LN5— responses increased sublinearly with pulse number because responses to successive pulses were progressively suppressed by response adaptation (Clemens et al., 2021) (Fig. 3A–C). This changed in the coincidence detector LN3, where the response to two pulses became supralinear. Here, the delayed rebound triggered by the first pulse coincided with direct excitation from the second pulse, producing a disproportionately strong response (Fig. 3D, E). This enhancement was further amplified downstream in LN4. For chirps containing still more pulses, the increase became sublinear, but responses remained elevated relative to stages upstream of coincidence detection. Removing coincidence detection by ablating the AN1 input to LN3 abolished the supralinear increase (Fig. 3B), confirming that it was generated by coincidence of the two inputs within LN3. Coincidence detection in LN3 thus produces a nonlinear increase in network activity at the behavioral threshold of two pulses per chirp. The minimum chirp duration therefore emerges from the same coincidence-detection mechanism that underlies pulse recognition, rather than from a dedicated chirp-specific computation (Fig. 3F).

**Figure 3.**
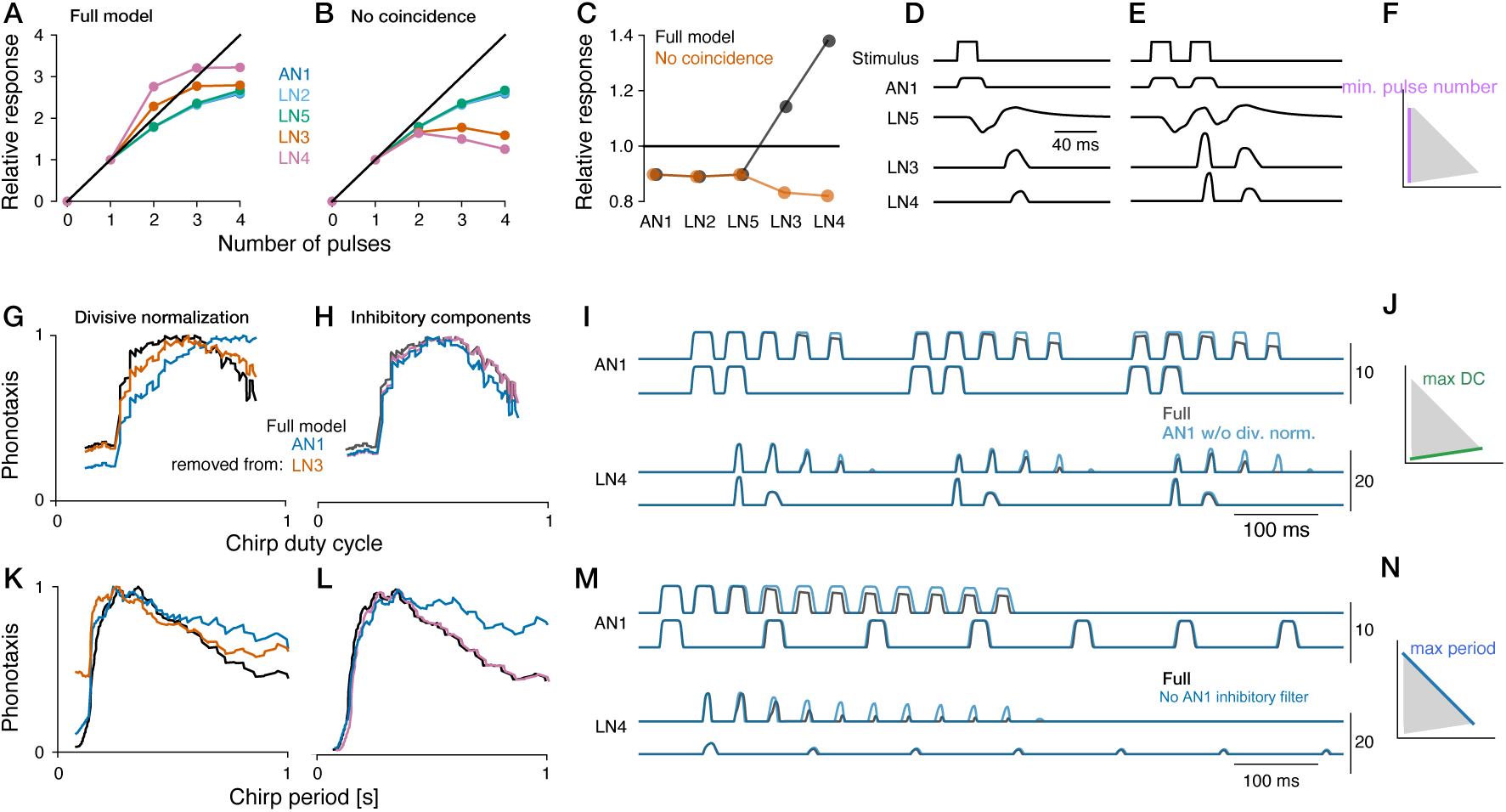
Coincidence detection and peripheral suppression generate chirp-selective boundaries. A,B. Response relative to response to a single-pulse stimulus across increasing pulse number at successive network stages in the full model (A) and after removing the direct AN1 input to LN3 (B). Neuron colors are AN1, blue; LN2, sky blue; LN5, bluish green; LN3, orange; and LN4, purple. The diagonal shows linear summation. Supralinear responses in the full model with intact coincidence detection create a behavioral boundary at the transition from one-to two-pulse stimuli. **C** Two-pulse response relative to the linear prediction from the one-pulse response in the full model (gray) and without coincidence detection (orange). Values above one indicate supralinear summation. **D,E** Stimulus and model responses to one-pulse (D) and two-pulse (E) chirps. **F** Coincidence detection establishes the minimum pulse-number boundary (purple). **G,H** Chirp-duty-cycle tuning in the full model (black) and after removing divisive normalization in AN1 (blue) or LN3 (orange; G), or the inhibitory component of the AN1 temporal filter (blue) or inhibition onto LN4 (purple; H). Removing divisive normalization in AN1 has the strongest effect on duty-cycle tuning. **I** AN1 (top) and LN4 (bottom) responses to three duty-cycle stimuli in the full model (gray) and after removing AN1 divisive normalization (blue). Perturbation colors identify the manipulated neuron, including when its effect is shown in a downstream neuron. Without divisive normalization in AN1, responses decay less strongly during a chirp in AN1 and downstream in LN4. **J** Peripheral suppression establishes the maximum duty-cycle boundary (green). **K,L** Chirp-period tuning in the full model (black) and after removing divisive normalization in AN1 (blue) or LN3 (orange; K), or the inhibitory component of the AN1 temporal filter (blue) or inhibition onto LN4 (purple; L). Removing the inhibitory filter component in AN1 has the strongest effect on chirp period tuning. **M** AN1 (top) and LN4 (bottom) responses to three chirp-period stimuli in the full model (gray) and after removing the inhibitory component of the AN1 temporal filter (blue). Removing the inhibitory filter component from AN1 reduces the decline in responses during longer chirps. **N** Peripheral suppression establishes the maximum chirp-period boundary (blue).

### Divisive normalization reduces responses to high chirp duty cycles

The lower duty-cycle boundary follows from the requirement for a minimum number of pulses per chirp. The decline in responses at high duty cycles, however, points to a different mechanism. High-duty-cycle songs contain more pulses per chirp and more overall sound energy, yet evoke weaker behavioral and model responses than songs with intermediate duty cycles. This non-monotonic relationship suggests that suppressive computations counteract the additional excitation provided by these extra pulses. We therefore asked which suppressive operations in the model generate the reduction in responses at high duty cycles. The model contains divisive normalization in AN1 and LN3, an inhibitory component in the AN1 temporal filter, and inhibition from LN2 onto LN4.

Of these suppressive operations, only removing divisive normalization in AN1 largely abolished the decline at high duty cycles, converting the band-pass duty-cycle tuning into an approximately monotonic increase with the number of pulses per chirp (Fig. 3G, H). By contrast, removing divisive normalization in LN3 or subtractive inhibition in AN1 and LN4 produced at most modest increases in responses at high duty cycles. In the intact model, AN1 normalization progressively attenuated responses to successive pulses, so that later pulses contributed increasingly little to the network output (Fig. 3I). Without normalization, successive pulses contributed more uniformly. Thus, divisive normalization in AN1, at the earliest stage of the pulse recognition network, is the principal mechanism generating the upper duty-cycle boundary (Fig. 3J).

### Peripheral suppression generates the upper chirp-period boundary

Coincidence detection explains the minimum chirp duration, whereas suppression in AN1 explains the decline in responses at high duty cycles. We next asked whether suppression also accounts for the upper chirp-period boundary. At constant duty cycle, increasing chirp period lengthens each chirp and therefore increases the number of pulses within it. Longer chirps could consequently recruit progressively stronger suppression, reducing responses despite the additional pulses.

Perturbing the model supported this hypothesis. As for high duty cycles, ablating suppressive computations in AN1 had the strongest effects on responses to long chirp periods. Removing either divisive normalization or the inhibitory component of the AN1 temporal filter substantially reduced the decline at long periods (Fig. 3K–M). Removing the inhibitory component of the temporal filter produced the larger effect, followed by divisive normalization in AN1, whereas normalization in LN3 and inhibition onto LN4 contributed comparatively little. The upper chirp-period boundary is therefore generated primarily by two suppressive computations in AN1. As chirps lengthen, these mechanisms progressively attenuate responses to successive pulses and thereby limit responses to songs with long chirp periods (Fig. 3M, N).

Together, all three behavioral boundaries arise from two computational principles already present in the pulse recognition network: coincidence detection establishes the minimum chirp duration, whereas peripheral suppression generates the decline at high duty cycles and the upper chirp-period boundary.

### The computations underlying pulse and chirp recognition are distributed differently across the network

The preceding analyses identified the computations responsible for each of the three behavioral boundaries. We next asked how pulse and chirp selectivity are organized across the network more generally. The modeled recognition network is anatomically largely serial (Fig. 1F) and contains no separate pulse-and chirp-processing branches. A conventional serial model would instead predict that short-timescale pulse features are computed first and then integrated downstream to generate longer-timescale chirp selectivity (Fig. 1C). To test this prediction, we quantified the overall effect of selectively removing each computation on the complete pulse-and chirp-tuning response fields (Fig. 4A–L).

**Figure 4.**
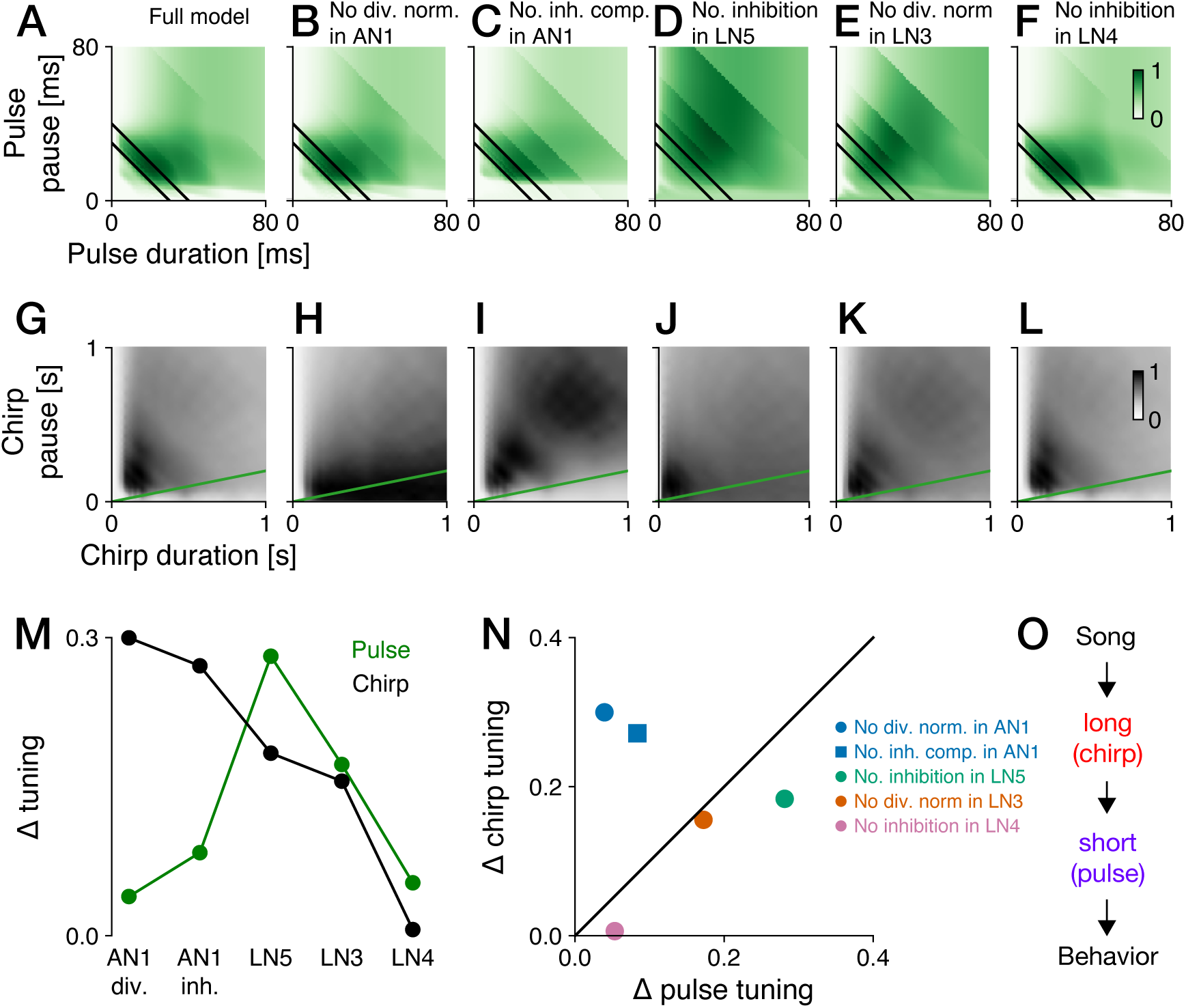
Suppressive computations contribute differently to pulse and chirp tuning in *G. bimaculatus*. A–F. Pulse preference fields for the full model (A) and after removing AN1 divisive normalization (B), the inhibitory filter component in AN1 (C), rebound-generating inhibition in LN5 (D), LN3 divisive normalization (E), or inhibition onto LN4 (F). Black lines mark the preferred pulse-period band in the full model for reference. **G–L** Corresponding chirp preference fields. Green lines mark the maximum duty-cycle boundary for the measured behavioral response. **M** Root-mean-square change in pulse (green) and chirp (black) tuning caused by perturbations at successive network stages. **N** Change in chirp tuning plotted against the corresponding change in pulse tuning for each perturbation. Colors identify the manipulated neuron: AN1, blue; LN5, bluish green; LN3, orange; and LN4, purple, as in Fig. 3. Blue circles indicate removal of AN1 divisive normalization and blue squares removal of the AN1 inhibitory filter. Points above the diagonal affect chirp tuning more strongly than pulse tuning. **O** Summary of the temporally interleaved organization: slow chirp selectivity arises early in the serial pathway (AN1), whereas fast pulse selectivity is completed downstream through rebound generation in LN5 and coincidence detection in LN3.

This analysis revealed an orderly spatial organization of temporal processing within the recognition network (Fig. 4M,N). Suppressive computations in the ascending auditory neuron AN1 had by far the largest effects on chirp tuning, consistent with their dominant role in establishing duty-cycle selectivity and the upper chirp-period boundary (Fig. 3G–N). Rebound generation in LN5 showed the opposite pattern, with a much larger effect on pulse tuning than on chirp tuning. LN3 divisive normalization had intermediate and similar effects on both temporal scales, whereas inhibition onto LN4 had small effects, particularly on chirp tuning.

Thus, although the network is anatomically largely serial, its functional organization does not follow a simple short-timescale-first, long-timescale-later hierarchy (Fig. 1C). The principal computations underlying chirp selectivity are already established in the ascending auditory pathway, whereas pulse selectivity continues to emerge downstream through rebound generation and coincidence detection (Fig. 4O). The dominant computations for the longer temporal scale therefore arise earlier in the network than those for the shorter temporal scale.

The concentration of chirp-selective computations in AN1 raises the possibility that variation in these early suppressive mechanisms could generate diversity in chirp preferences across species. We therefore next asked how changes in AN1 suppression reshape chirp tuning and whether they can reproduce the known diversity of chirp preferences across species.

### Variation in peripheral suppressive computations reproduces species differences in chirp preferences

We systematically varied the strength of divisive normalization and the inhibitory filter in AN1 while leaving all other model parameters unchanged. The two suppressive computations controlled complementary aspects of chirp preference. Reducing divisive normalization progressively transformed duty-cycle tuning from narrow band-pass, through broad band-pass, to high-pass tuning with a preferred duty cycle approaching one (Fig. 5A). Chirp-period tuning was affected comparatively little, exhibiting only a modest increase in responses to long chirp periods (Fig. 5B). In contrast, weakening the inhibitory filter had relatively little influence on duty-cycle tuning but substantially broadened chirp-period tuning by increasing responses to long chirp periods (Fig. 5C,D). Together, these two suppressive computations control two principal dimensions of chirp preference (Fig. 5E,F).

**Figure 5.**
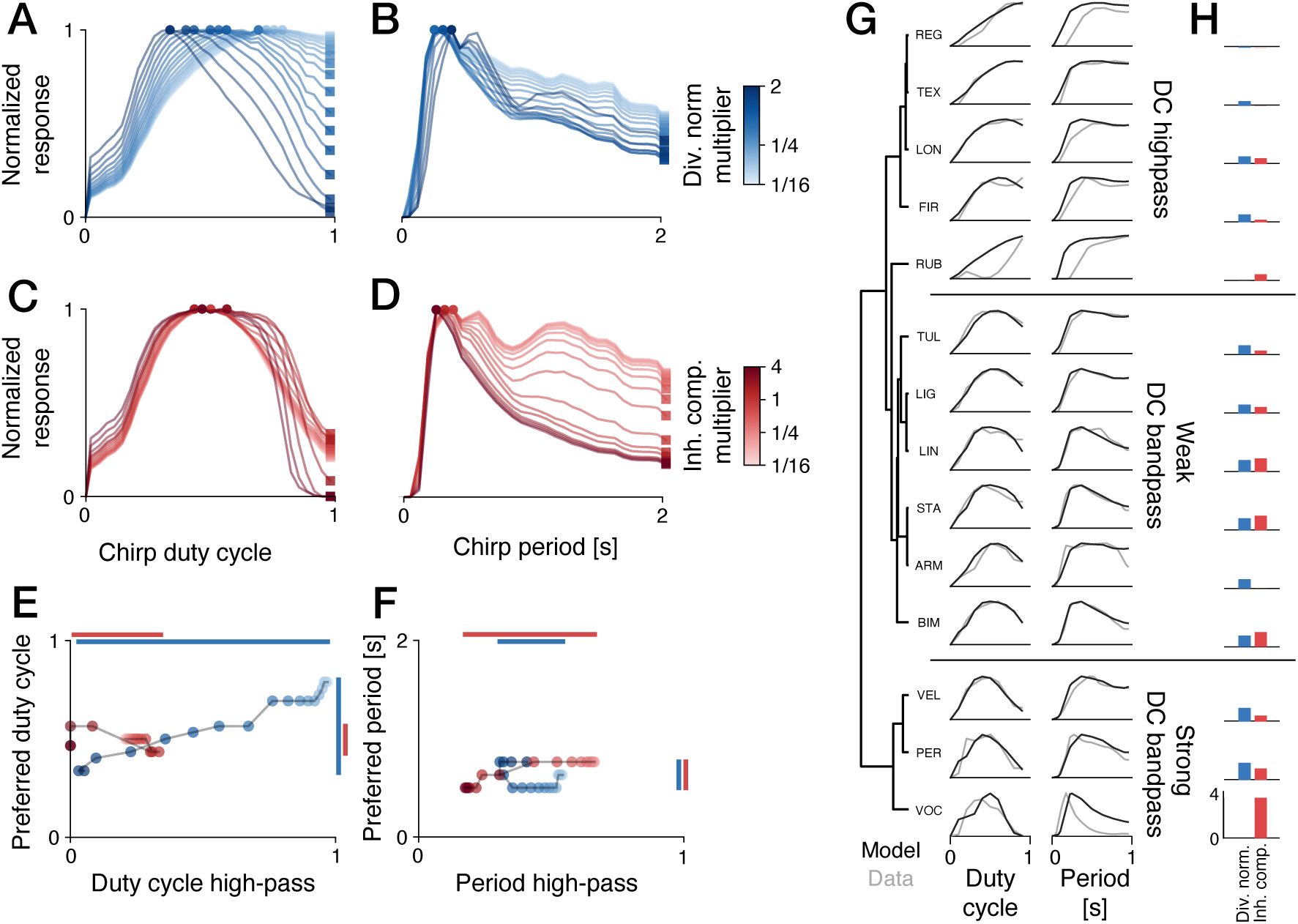
Variation in peripheral suppression reproduces the dominant diversity of chirp tuning across species. A,B. Maximum-normalized chirp-duty-cycle (A) and chirp-period (B) tuning as AN1 divisive-normalization strength is varied. Blue shade indicates the parameter multiplier (see color bar in B). Circles and squares indicate the responses at the preferred and largest sampled values, respectively, for each tuning curve. **C,D** Corresponding tuning as the inhibitory component of the AN1 temporal filter is varied. Red shade indicates the parameter multiplier (see color bar in D). Markers have the same meaning as in A,B. **E,F** Preferred chirp duty cycle versus the duty-cycle high-pass index (E), and preferred chirp period versus the period high-pass index (F), for the individual parameter sweeps. Same color-coding as in A–D. Vertical and horizontal lines indicate the range of values covered by the two parameter sweeps, with AN1 divisive normalization more strongly affecting duty-cycle tuning (E, blue), and the AN1 inhibitory component more strongly affecting period tuning (F, red). The duty-cycle (E) and period (F) high-pass indices are the responses at the largest sampled duty cycle and chirp period, respectively, divided by the corresponding peak response. **G** Best-fitting model (black) and behavioral (gray) tuning for chirp duty cycle (left) and period (right). Species are ordered from high-pass tuning at the top through weak to strong duty-cycle band-pass tuning at the bottom. The dendrogram shows similarity in duty-cycle tuning, and labels identify the three groups. See Table S2 for full species names. **H** Divisive-normalization (blue) and inhibitory-filter (red) multipliers of the best-fitting model for each species. Bars indicate the fold-change (0–4) of the respective parameters relative to the corresponding parameter of the stabilized reference model.

We next asked how much of the diversity of chirp preferences observed across species could be accounted for by varying these two suppressive computations. To this end, we compared two-dimensional chirp preference fields from *G. bimaculatus* and 13 additional cricket species. Their phylogenetic relationships are shown in Supplementary Fig. S4. The complete chirp preference fields contained substantial species-specific structure (Supp. Fig. S5). To reduce this complexity, we focused on two one-dimensional tuning profiles: chirp-duty-cycle and chirp-period tuning (Fig. 5G). Along these profiles, species differences were dominated by relatively simple variation between high-pass and band-pass response shapes. Note that this reduction captures major axes of variation, but not the full species-specific structure of each preference field. Species ranged from high-pass duty-cycle tuning, with strong responses to continuous pulse trains, to increasingly narrow band-pass tuning that completely rejected continuous pulse trains. Likewise, chirp-period tuning ranged from broad high-pass tuning to narrow band-pass tuning with weak responses to long chirp periods. High-pass chirp-duty-cycle tuning was consistently associated with high-pass chirp-period tuning, suggesting that weak suppression can contribute to both response properties. In contrast, species with broad or narrow band-pass duty-cycle tuning exhibited substantial diversity in chirp-period tuning, indicating that the two tuning dimensions can be adjusted independently.

We therefore asked whether jointly varying divisive normalization and the inhibitory filter in AN1 could reproduce the tuning of individual species. Fitting the two parameters to each species produced model profiles that captured the principal combinations of duty-cycle and chirp-period tuning for most species (Fig. 5G), using species-specific combinations of the two suppressive computations (Fig. 5H). The tuning of *G. vocalis*, *G. rubens* and *G. regularis* was reproduced qualitatively, although not in every detail. *G. vocalis* responses declined sharply at long chirp periods, more abruptly than predicted by the model. In *G. rubens* and *G. regularis*, model responses rose at shorter chirp periods than the behavioral responses. In addition, the model overestimated responses to low and intermediate duty cycles in *G. rubens*. This indicates that additional nonlinearities or computations contribute to chirp selectivity in these species. The relationship between species-specific chirp tuning and the fitted parameters was consistent with the sweeps of individual parameters (Fig. S7). Species with weaker normalization preferred higher duty cycles (Spearman *ρ* = *−0.60*, *p* = *0.02*), and species with a weaker inhibitory filter had higher period high-pass indices (*ρ* = *−0.89*, *p* = *2 × 10 ^−5^*) and preferred longer chirp periods (*ρ* = *−0.79*, *p* = *9 × 10 ^−4^*).

Together, these results suggest that quantitative changes in two suppressive computations can reproduce major cross-species differences along the chirp-duty-cycle and chirp-period tuning profiles. They do not account for all structure in the complete preference fields, but show that prominent high-pass-to-band-pass variation can arise through retuning of early sensory computations.

## Discussion

The usual distinction between serial and parallel recognition assumes that temporal scale maps onto circuit organization. In a classical parallel architecture, separate channels analyze different timescales before their outputs converge. In a conventional serial hierarchy, short elements are recognized first and then assembled into progressively longer units (Fig. 1C). Cricket behavior and the independently constrained circuit model support a serial organization with an inverted temporal hierarchy: in the functionally and anatomically largely serial recognition network (Fig. 1F), suppressive mechanisms near the circuit input establish selectivity for long-timescale chirp features, whereas selectivity for the faster pulse pattern is completed downstream (Figs. 3G–N and 4M–O). Serial wiring therefore does not require computation to proceed from short to long timescales.

This organization provides a circuit interpretation of earlier descriptions of pulse-and chirp-scale preferences as a weighted AND-like operation (Grobe et al., 2012; Meckenhäuser et al., 2013). In the serial architecture proposed here, mistimed pulses weaken coincidence detection, whereas unattractive chirp structure suppresses the gain of the signal passed downstream. Selectivity at either scale can therefore reduce the output below the response threshold, while favorable structure at one scale compensates only partly for suboptimal structure at the other.

This organization is possible because AN1 carries information about both timescales in different aspects of its response. Pulse timing remains encoded in the temporal pattern of pulse-evoked activity, whereas divisive normalization and the inhibitory temporal filter convert recent pulse history into changes in response gain (Fig. 3). In AN1, the effect of suppression on any single pulse is modest, but small gain differences propagate through the circuit and are amplified by downstream nonlinearities, like thresholding, generating selectivity for chirp duty cycle and period in the output neuron of the network, LN4 (Fig. 3I, M). In parallel, pulse timing is converted into firing rate through the LN5–LN3 rebound and coincidence-detection mechanism which is sensitive to pulse timing and relatively robust to gain (Schöneich et al., 2015) (Fig. 3A–F). LN4 combines these signals, so its firing rate encodes both short-timescale pulse structure and long-timescale chirp structure and reproduces much of the behavioral selectivity at both scales (Fig. 2G–M, 4A, G). Long patterns can thus be recognized through the internal state of a circuit without an explicit representation of the chirp as a discrete object.

This division of labor explains how pulse and chirp preferences can vary partly independently without separate anatomical modules. Chirp history and pulse structure are carried primarily by gain and timing, respectively, and are most sensitive to different computations in different neurons (Fig. 4M, N)—suppressive computations in AN1 (Fig. 3G–N) and rebound generation in LN5 and coincidence detection in LN3 (Fig. 3A–F). Behavioral independence between features therefore need not imply independent processing channels.

The cross-species analysis suggests that this organization also provides a simple route for evolutionary diversification. Variation in two suppressive computations in AN1 reproduces major high-pass-to-band-pass differences among species along the chirp-duty-cycle and chirp-period tuning profiles (Fig. 5A–H). Divisive normalization primarily changes tolerance for chirps with large duty cycles, whereas the inhibitory temporal filter more strongly changes tolerance for long chirp periods. Quantitative changes to existing sensory computations may therefore shift preferences along major axes of natural song variation without requiring the evolution of a new chirp-recognition pathway. Because AN1 supplies the entire downstream network, however, such changes are unlikely to be free of pleiotropic effects. The relative preservation of pulse tuning in the model suggests that downstream rebound and coincidence-detection mechanisms buffer some of these effects, but the extent of this buffering remains an empirical question. Of special interest for future study are species with unusual songs, for instance, *G. bryanti*, whose calling songs usually consist of single-pulse chirps with a period of about 1 s and for which a coincidence mechanism is unlikely (Weissman et al., 2019).

These conclusions establish computational sufficiency rather than a complete biological mechanism. The model was constrained primarily by pulse-scale electrophysiology and behavior and required a reduction in AN1 normalization to remain stable during long stimulus sequences. The model places the suppressive computations at the AN1 stage, before downstream pulse recognition, but does not establish whether its biological source lies within AN1 or is inherited from upstream neural inputs, like the auditory receptor or the omega neurons (Benda and Hennig, 2008; Clemens et al., 2015b). The model’s recovery of the principal chirp boundaries after this adjustment shows that a dedicated chirp stage is not required to explain the main behavioral selectivity, but does not exclude additional chirp-sensitive computations elsewhere in the auditory pathway. Additional computations likely further reduce responses to long chirp pauses, which the model overestimated (Fig. 2J). The model also failed to reproduce some structure in the chirp-duty-cycle and period tuning curves of some species (Fig. 5). These discrepancies may reflect species differences in computations outside AN1, interactions among song features not captured by the two summary axes, or limitations of the current model.

The proposed organization nevertheless makes direct experimental predictions. The strength and recovery time of suppression in AN1 should predict an individual’s or species’ rejection of high-duty-cycle and long-period chirps. Weakening this suppression should shift the corresponding upper chirp boundaries more strongly than pulse preference, whereas perturbing rebound generation or coincidence detection should have larger effects on pulse tuning and the minimum number of pulses required per chirp. Broad chirp selectivity should therefore already be predictable in AN1 responses before sharp pulse selectivity emerges downstream. Recordings from identified neurons in the network across species would test these predictions directly.

Other compact auditory systems illustrate contrasting solutions to the same multiscale problem. Frogs follow the classical short-first, long-later hierarchy: interval-counting neurons serially accumulate correctly timed short intervals into a longer-timescale response (Alder and Rose, 1998; Edwards et al., 2002; Edwards et al., 2007; Rose et al., 2015). Grasshoppers instead process syllable duration, pause duration and syllable number in parallel channels that subsequently converge (Creutzig et al., 2009; Creutzig et al., 2010). *Drosophila* may resemble crickets in temporal order, because response adaptation in early auditory neurons encodes the components from which song-bout duration can be recovered before selectivity for the much shorter interpulse interval emerges downstream (Clemens et al., 2015a; Deutsch et al., 2019; Wang et al., 2021). The fly auditory system is highly parallel, however (Baker et al., 2022; Schlegel et al., 2024; Dorkenwald et al., 2024), so this order may arise across distinct processing branches rather than through temporally interleaved computations within a single serial pathway.

These comparisons also qualify a common interpretation of cortical temporal hierarchies. Progressively longer temporal receptive windows and intrinsic timescales along a cortical hierarchy are often taken to imply that long-timescale features are constructed downstream from completed short-feature representations (Kiebel et al., 2008; Hasson et al., 2008; Lerner et al., 2011; Honey et al., 2012; Murray et al., 2014; Chaudhuri et al., 2015; Hasson et al., 2015; Runyan et al., 2017). The cricket circuit shows that anatomical order, integration timescale and behavioral selectivity need not covary. A slowly recovering process near the sensory input can establish a long-timescale boundary while later stages are still computing selectivity for much faster structure.

More generally, multiscale recognition requires neither a separate module for every timescale nor a hierarchy ordered from short to long. A compact circuit can use its internal state to represent several timescales simultaneously and downstream nonlinearities to read out different aspects of that state. Such temporally interleaved computation provides an economical and evolvable solution for recognizing hierarchical signals.

## Methods

### Behavioral data and acoustic stimuli

For *G. bimaculatus*, we analyzed a chirp preference data set comprising 60 synthetic songs distributed across chirp duration and chirp pause (Fig. 2A). See Grobe et al. (2012) for experimental details. The stimulus grid was sampled most densely around the approximately 400-ms chirp period of the conspecific song. Every chirp contained rectangular 20-ms pulses separated by 20-ms pulse pauses, thereby holding the pulse duty cycle at 0.5 while chirp duration and pause varied. Chirp period was defined as chirp duration plus chirp pause, and chirp duty cycle as chirp duration divided by chirp period. The measured phonotaxis scores were interpolated onto the regular duration-pause grid used for visualization and model comparison and were normalized to the maximum response within each field. For visualization, negative response values were clipped to zero and two zero-response anchors were added at chirp duration–pause coordinates (0, 0) and (0, 1000) ms. The data were interpolated by natural-neighbor interpolation, requiring at least four neighbors, onto a regular 5-ms duration–pause grid spanning 0–1000 ms. Duplicate coordinates were separated by sub-0.01-ms numerical jitter before interpolation to avoid numerical errors, and negative interpolated values were set to zero.

One-dimensional response profiles were derived from the measured data within fixed bands of the two-dimensional field selected to encompass the response peaks. Chirp-duration and pulse-number profiles used chirp pauses of 160–240 ms; the chirp-pause profile used chirp durations of 120–200 ms; the chirp-period profile used duty cycles of 0.4–0.6; and the chirp-duty-cycle profile used chirp periods of 240–320 ms. When multiple stimuli had the same value on the profile axis, their maximum response was retained. Profiles were sorted and smoothed with a three-point running mean after padding the endpoints with their adjacent values. Behavioral and model profiles were normalized by the respective maximum response across the full stimulus sets. The three boundaries were used as descriptive summaries rather than fitted inferential thresholds: minimum chirp duration of approximately 40 ms, chirp periods between approximately 150 and 600 ms, and maximum chirp duty cycle of approximately 0.8. They therefore summarize the topology of the measured response field and are not estimates of categorical psychophysical thresholds.

### Comparative chirp preference data

The comparative analysis included *G. bimaculatus* and 13 additional cricket species (Fig. S5). Chirp data for nine of the 14 species, including *G. bimaculatus*, were published (Grobe et al., 2012; Hennig et al., 2016; Blankers et al., 2015; Gray et al., 2016). The chirp data for the remaining five species are unpublished (Table S2). Behavioral experiments were performed as for *G. bimaculatus*. Each field was interpolated by natural-neighbor interpolation onto a regular 20-ms duration–pause grid spanning zero to the largest duration or pause sampled for that species. For phylogenetic context, we pruned the multilocus species tree of Gray et al. (2020) to these 14 species (Supplementary Fig. S4), using its Florida *G. firmus* sample to represent the source population of the behavioral data (Gray et al., 2016). Chirp-duty-cycle profiles were calculated over species-specific chirp-period windows selected to encompass the species-specific response peak (see Table S2). Chirp-period profiles were calculated for duty cycles of 0.33– 0.67, except in *G. rubens*, for which duty cycles of 0.9–1.0 were used because responses were concentrated at high duty cycles. Within each band, duplicate values on the profile axis were represented by their maximum response and then averaged within 15 quantile-spaced bins. A leading zero-response anchor was retained at half the smallest supported value on the profile axis, and each profile was normalized to its maximum. Preferred chirp duty cycle or period was the center of the bin with the maximum response. The period and duty-cycle high-pass indices were the response to the largest period or duty-cycle value divided by the maximum response. The index therefore ranged from zero for rejection of the endpoint stimulus to one when the response remained maximal at the endpoint. Species in Fig. 5G were ordered by average-linkage hierarchical clustering of the Euclidean distances between the final two points of their normalized duty-cycle profiles, using optimal leaf ordering.

### Pulse-recognition network model and long-stimulus stabilization

We used the mechanistic model of the identified *G. bimaculatus* pulse-recognition network described previously (Clemens et al., 2021), implemented in Python with JAX. The model represents the ascending auditory neuron AN1 and local neurons LN2, LN3, LN4 and LN5 using linear temporal filters, static nonlinearities, synaptic delays and gains, post-inhibitory rebound, coincidence detection, divisive normalization and inhibitory interactions. The published base parameters were determined from pulse-scale electrophysiology and behavior. The published model was developed for relatively short pulse stimuli. With the longer sequences used here, it produced a slow oscillation in response magnitude and non-smooth response fields. We therefore rectified the AN1 signal before divisive normalization, replacing *x* (*t*) at that stage with max[*0*, *x* (*t*)], and reduced the AN1 normalization weight from 10 to 4. These changes stabilized the long-duration response while leaving the qualitative pulse and chirp tuning largely unchanged (Supplementary Fig. S3).

The model was evaluated at 1 kHz. Acoustic stimuli were unit-amplitude rectangular pulse trains. Chirps contained complete 20-ms pulses and 20-ms pulse pauses; chirp duration was converted to an integer number of complete pulse periods, the resulting chirp was followed by the specified chirp pause, and complete chirps were repeated for the requested song duration. Every stimulus began with 50 ms of silence and any unused interval after the last complete chirp was zero-padded. All analyses used 10-s songs, except for the long-stimulus stability analysis, which used 50-s songs. All species models retained 20-ms pulse durations and pauses.

For the main model–behavior comparison and chirp perturbation fields, chirp duration and pause each took 32 geometrically spaced values from 20 to 1020 ms. Species sweeps and fits used 32 values from 40 to 1040 ms. The model-stabilization comparison (Fig. S3) used 64 values from 20 to 1000 ms and 50-s songs. Pulse response fields used all integer pulse durations and pauses from 0 to 79 ms in 1-ms steps, presented in chirps with 200-ms duration and 200-ms pause. Model fields were interpolated with a linear radial-basis function with smoothing parameter 20 onto regular 5-ms grids for the main and perturbation analyses and a regular 10-ms grid for the species analysis.

### Prediction of behavior from model responses and quantification of model fit

The firing rate of LN4 is linearly related to phonotaxis (Schöneich et al., 2015; Clemens et al., 2021). We therefore predicted phonotaxis, *p*, via the mean LN4 output, *r* (*t*), over all samples in the generated trace:

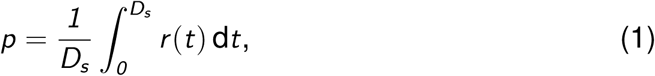

where *D_s_* includes the initial 50-ms silent interval. Model response fields were normalized to their maxima before comparison with behavior. To quantify model fit, behavioral values were linearly interpolated onto the model x-values over the behaviorally sampled range. Agreement between the interpolated behavioral values and corresponding model values was quantified as the mean-squared error. The separate objective and profile normalization for species fitting are described below.

### Computational perturbations and impact scores

We isolated individual computations by setting only the relevant model parameter to zero and leaving all other parameters fixed. Divisive normalization was removed jointly or selectively by setting the normalization weights in AN1 and/or LN3 to zero. The inhibitory component of the AN1 temporal filter was removed by setting its gain to zero. Rebound-generating inhibition in LN5 was removed by setting the duration of the differentiating LN5 filter to zero, and inhibition onto LN4 was removed by setting the LN2–LN4 synaptic gain to zero. Coincidence detection was disrupted by setting the direct AN1–LN3 synaptic gain to zero.

For the pulse-number analysis, chirp period was fixed at 320 ms and chirp duration was increased from 0 to 320 ms in 40-ms increments, producing chirps containing zero to eight pulses. Responses at each network stage were averaged over time. Supralinearity of the two-pulse response was quantified as *R_2_* /(*2 R_1_*), where *R_1_* and *R_2_* are responses to one-and two-pulse chirps; values above one indicate a response larger than the linear prediction.

For each perturbation, the complete pulse and chirp response fields were recomputed and separately maximum-normalized. The impact of perturbation *j* was quantified as

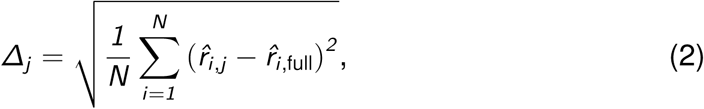

where *i* indexes the *N* stimulus-grid points and hats denote maximum-normalized responses. The calculation was applied separately to the pulse and chirp fields, allowing each computation to be placed in the pulse-impact versus chirp-impact plane in Fig. 4N.

### Cross-species parameter sweep

To test whether peripheral suppression could span the observed species diversity, we varied the strengths of AN1 divisive normalization and the inhibitory component of the AN1 temporal filter while keeping all other network parameters at their *G. bimaculatus* values. For the parameter sweeps in Fig. 5A–F, each parameter was varied separately at 19 logarithmically spaced multipliers: *2 ^−4^* –*2^1^* for divisive normalization and *2 ^−4^* –*2^2^* for the inhibitory filter. Each variant was evaluated on the 32-by-32 species chirp grid described above using 10-s songs to accommodate the longer chirp periods of some species. Duty-cycle profiles used chirp periods of 200–500 ms, and period profiles used duty cycles of 0.33–0.67. Profiles were normalized to their maxima after adding the leading zero-response anchor.

For model selection, duty-cycle curves were linearly interpolated at 10 values between 0 and 0.9. Chirp-period curves were interpolated at 40 values between 0 and 950 ms. Missing experimental bins were filled by linear interpolation over bin indices; interpolation onto the fitting grids held the endpoint response constant outside each profile’s sampled range. The best-matching joint model for each species minimized

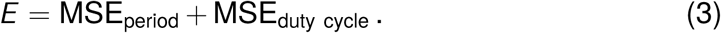

We optimized the two AN1 parameter multipliers for each species. The three sweep variants with the lowest values of *E* were used as independent starting points. From each starting point, *E* was minimized with the COBYQA, Powell, and Nelder–Mead algorithms, with both multipliers bounded between 0 and 4 and a limit of 200 model evaluations per run. We retained the lowest-error solution across the starting points and algorithms.

## Data and code availability

The original network model was published in (Clemens et al., 2021) and is deposited at https://github.com/janclemenslab/cricketnet. Behavioral data and code for all computational experiments and figures can be found at https://github.com/janclemenslab/cricket-chirps.

**Table 1.** Python packages used for data processing, model simulation, analysis and figure generation.

| Package | Reference |
| --- | --- |
| NumPy | (Harris et al., 2020) |
| pandas | (McKinney, 2010) |
| SciPy | (Virtanen et al., 2020) |
| JAX | (Bradbury et al., 2018) |
| MetPy | (May et al., 2022) |
| scikit-learn | (Pedregosa et al., 2011) |
| Matplotlib | (Hunter, 2007) |
| glasbey | (Glasbey et al., 2007) |

## Acknowledgements

We thank several students for help and support in conducting behavioral tests. This study was funded by a grant from the Deutsche Forschungsgemeinschaft (DFG, German Research Foundation) to MH and JC as part of the SPP 2205, project number 430158535.

## Supplementary Information

**Table S1.** Parameters of the calling song of *G. bimaculatus* males. Data from 93 males over 1 minute of song.

| Song Parameter | Mean±std | Range |
| --- | --- | --- |
| Pulse period | 40.8± 6.4 ms | 22.9-55.6 ms |
| Pulse rate | 24.5± 4.6 pulses/s | 17.9-43.6 pulses/s |
| Pulse DC | 29± 16% | 6-72% |
| Chirp period | 445.1±123.8 ms | 225.3-868.4 ms |
| Chirp rate | 2.2±.7 chirps/s | 1.2-4.4 chirps/s |
| Chirp DC | 30±10% | 10-50% |

**Table S2.**
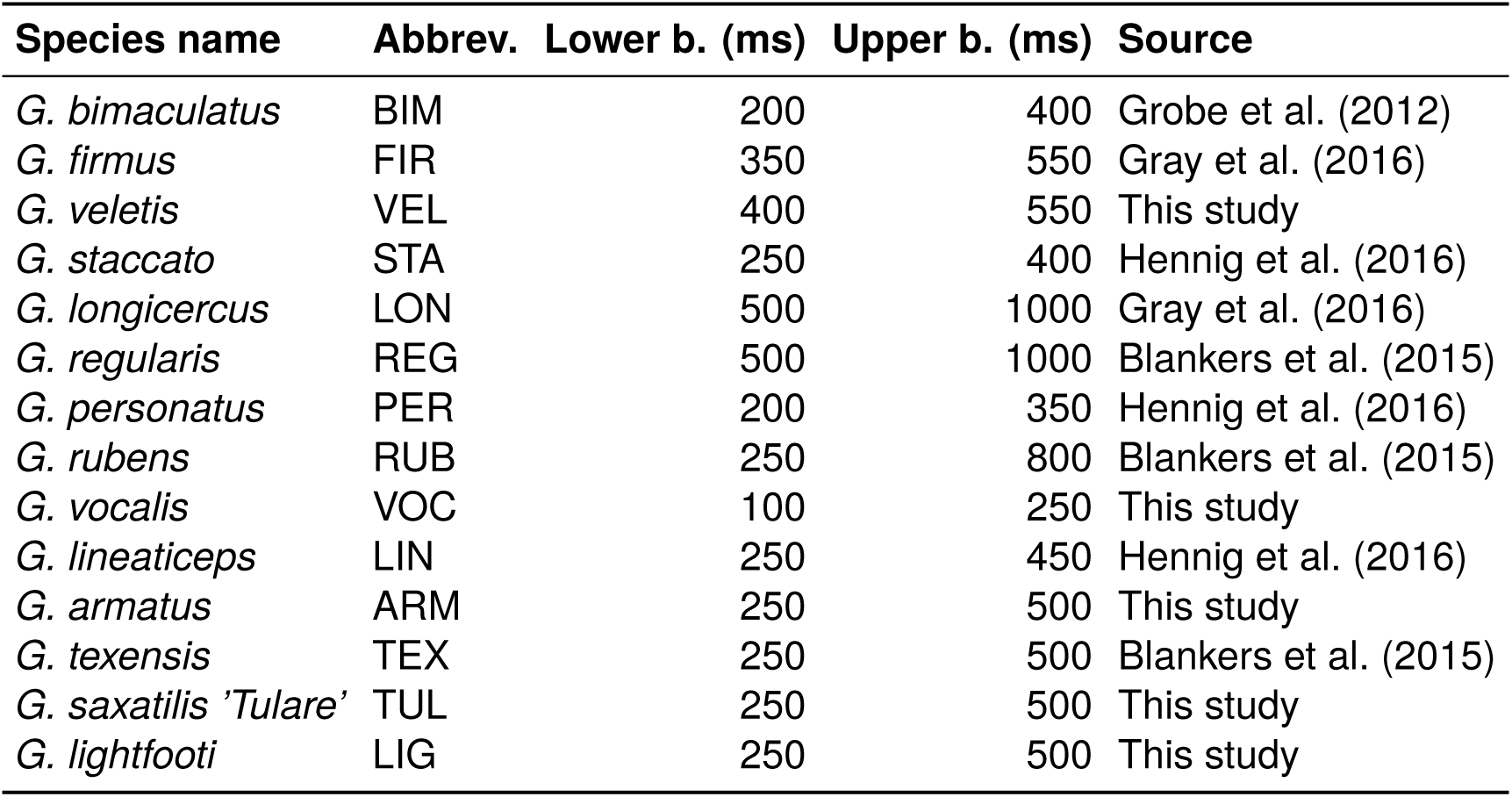
Chirp-period windows (lower and upper bound) used to calculate duty-cycle tuning for the 14 species with complete response fields. Species names according to Weissman and Gray (2019).

---

**Figure S1.**
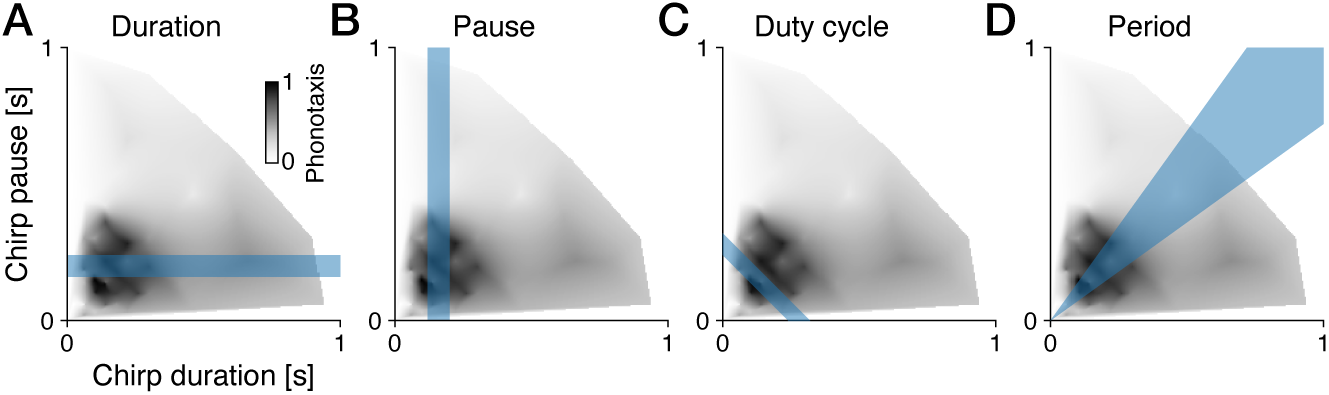
Parameter ranges for the four transects through the chirp response field. A–D. The parameter ranges (blue shaded area) over which the four transects (duration, A; pause, B; duty cycle, C; period, D) in Fig. 2 were calculated, overlaid on top of the chirp response field for *G. bimaculatus*.

**Figure S2.**
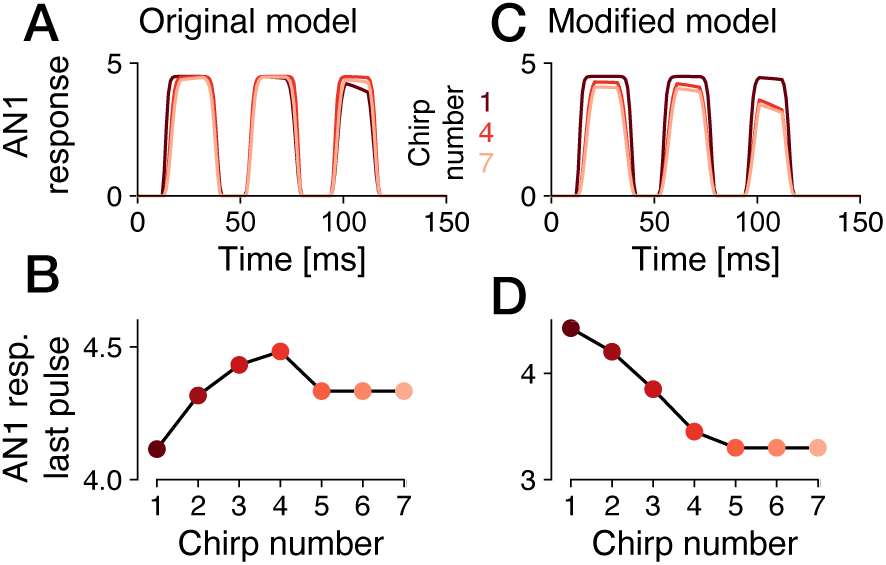
Long-timescale behavior in the original and the modified model. **A** Model AN1 responses (original parameters, Clemens et al. (2021)) to chirps 1, 4, and 7 (color coded, see legend) in a song. Response amplitude is lowest for the first chirp, peaks at chirp 4, and declines slightly afterwards. **B** Average model AN1 responses to the last pulse in a chirp as a function of chirp number in a song. **C, D** Same as A, B but with modified parameters. Responses now decay with chirp number and saturate after 4 chirps.

**Figure S3.**
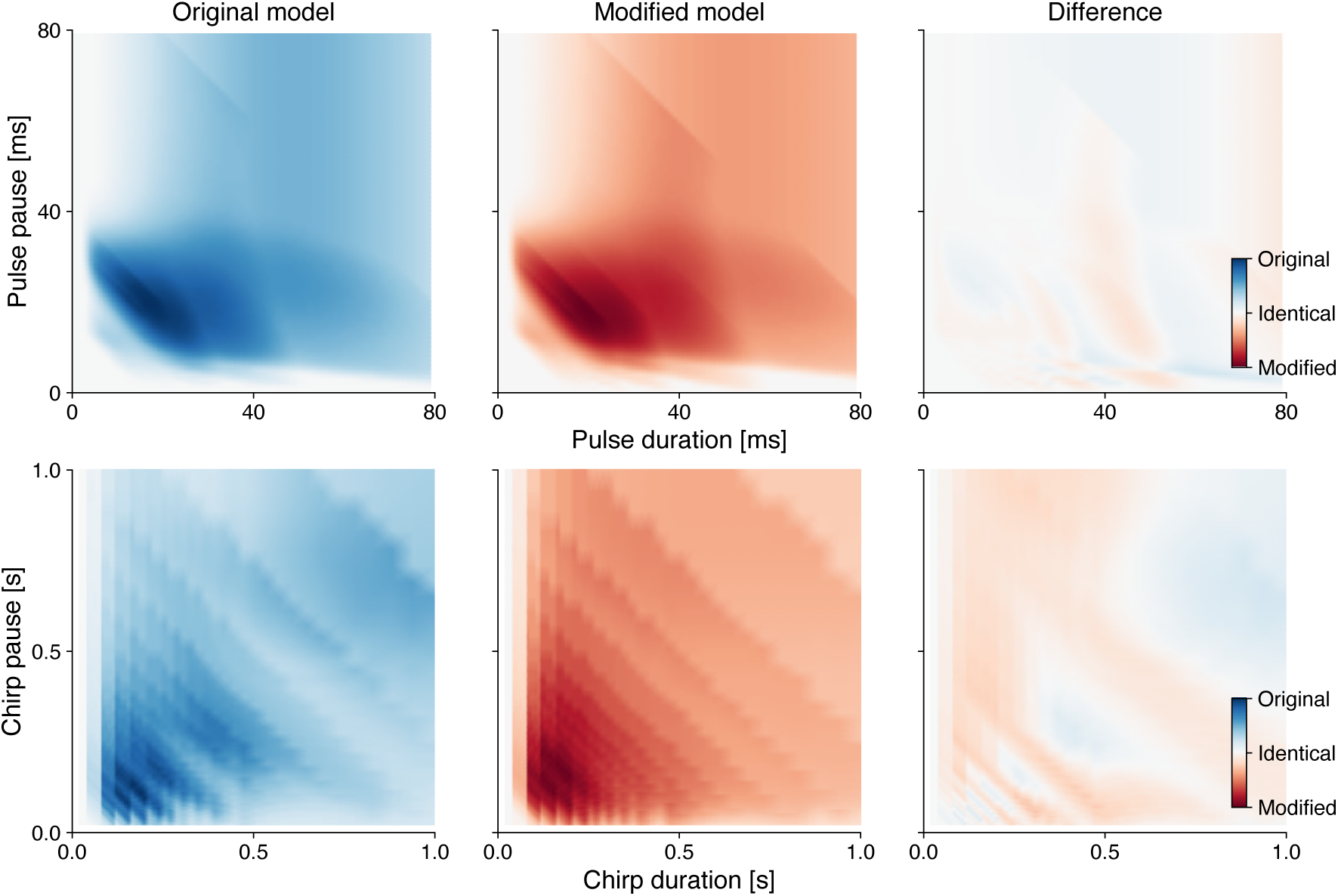
The long-stimulus model modification preserves pulse and chirp tuning. Pulse preference fields (top row) and chirp preference fields (bottom row) for the original model (left, blue), modified model (center, red), and their signed difference (right). Fields were maximum-normalized before subtraction. Blue indicates a larger response in the original model, red a larger response in the modified model, and white identical responses.

**Figure S4.**
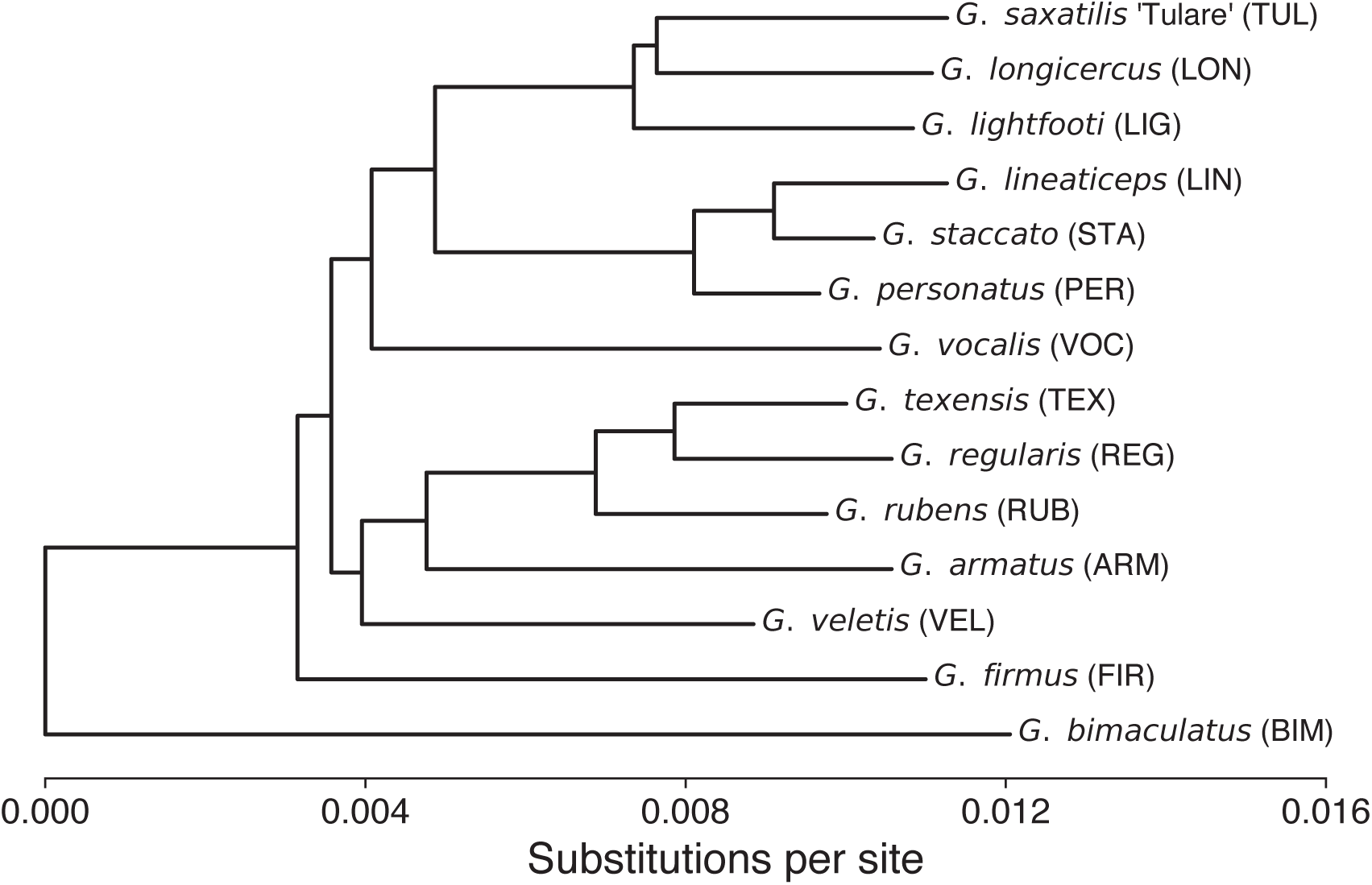
Phylogenetic relationships among the 14 cricket species in the comparative analysis. The multilocus species tree of Gray et al. (2020) was pruned to the species in Table S2. Branch lengths show substitutions per site. The Florida *G. firmus* sample was used to represent the source population of the behavioral data. The tree provides evolutionary context for the species comparison; no phylogenetically corrected analysis was performed.

**Figure S5.**
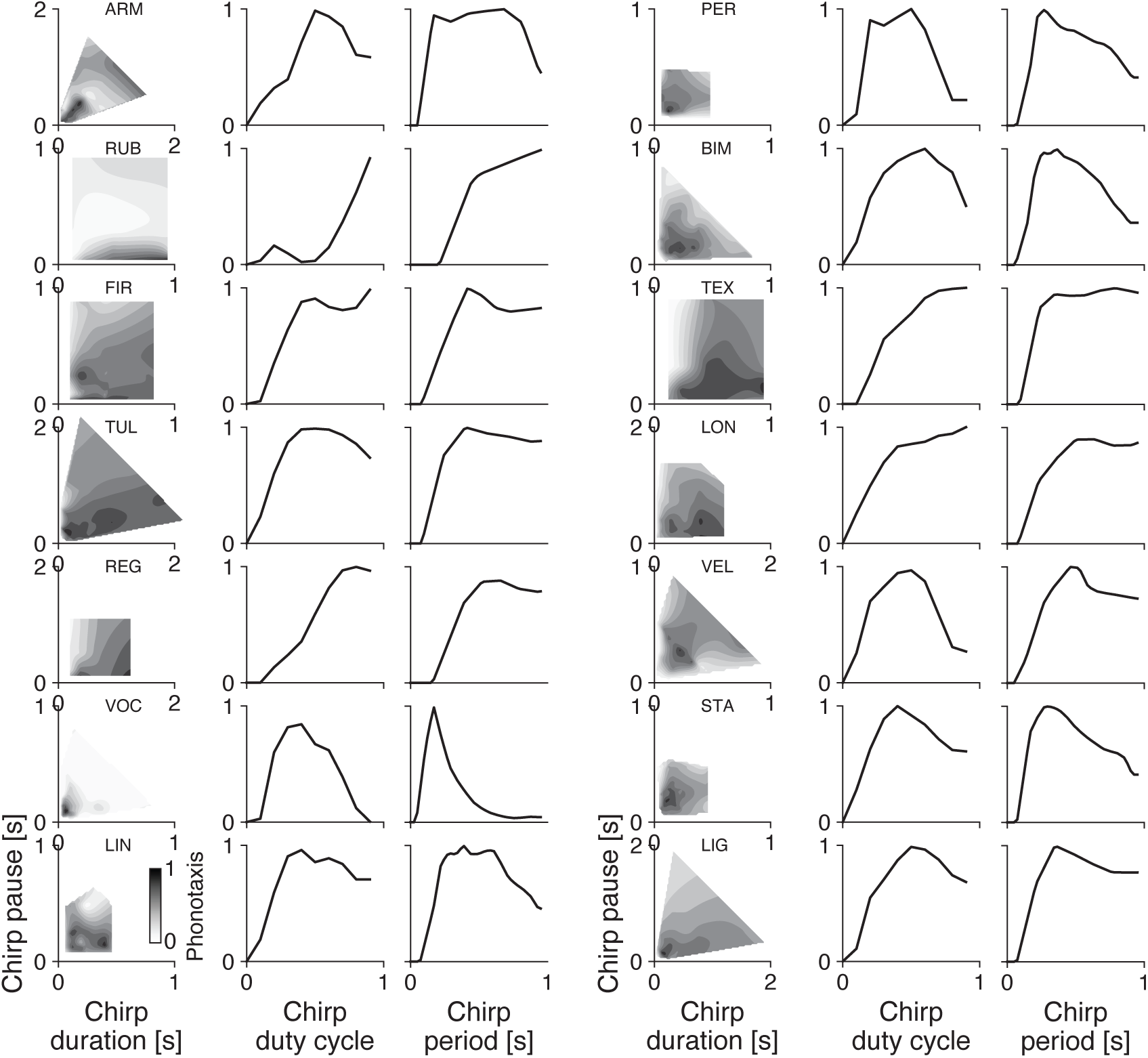
Chirp preference fields and one-dimensional tuning curves for 14 cricket species. For each species, the left panel shows the normalized chirp duration–pause response field, the center panel shows chirp-duty-cycle tuning within the species-specific chirp-period window in Table S2, and the right panel shows chirp-period tuning at duty cycles of 0.33–0.67 (0.9–1.0 for *G. rubens*). All tuning curves are normalized to their maxima. See Table S2 for full species names.

**Figure S6.**
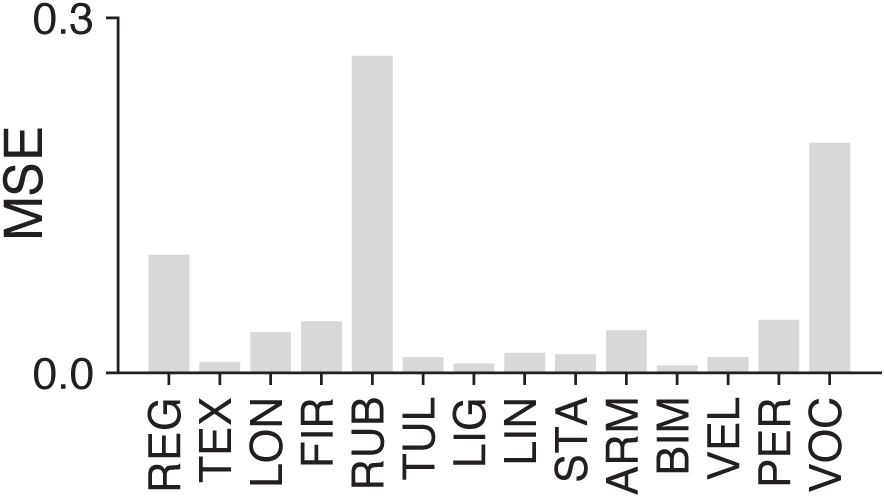
Model errors across species. Combined fitting error for the best-fitting model for each species. Error is the sum of the mean-squared differences between behavioral and model chirp-period and chirp-duty-cycle tuning curves. Lower values indicate better fits. See Table S2 for full species names.

**Figure S7.**
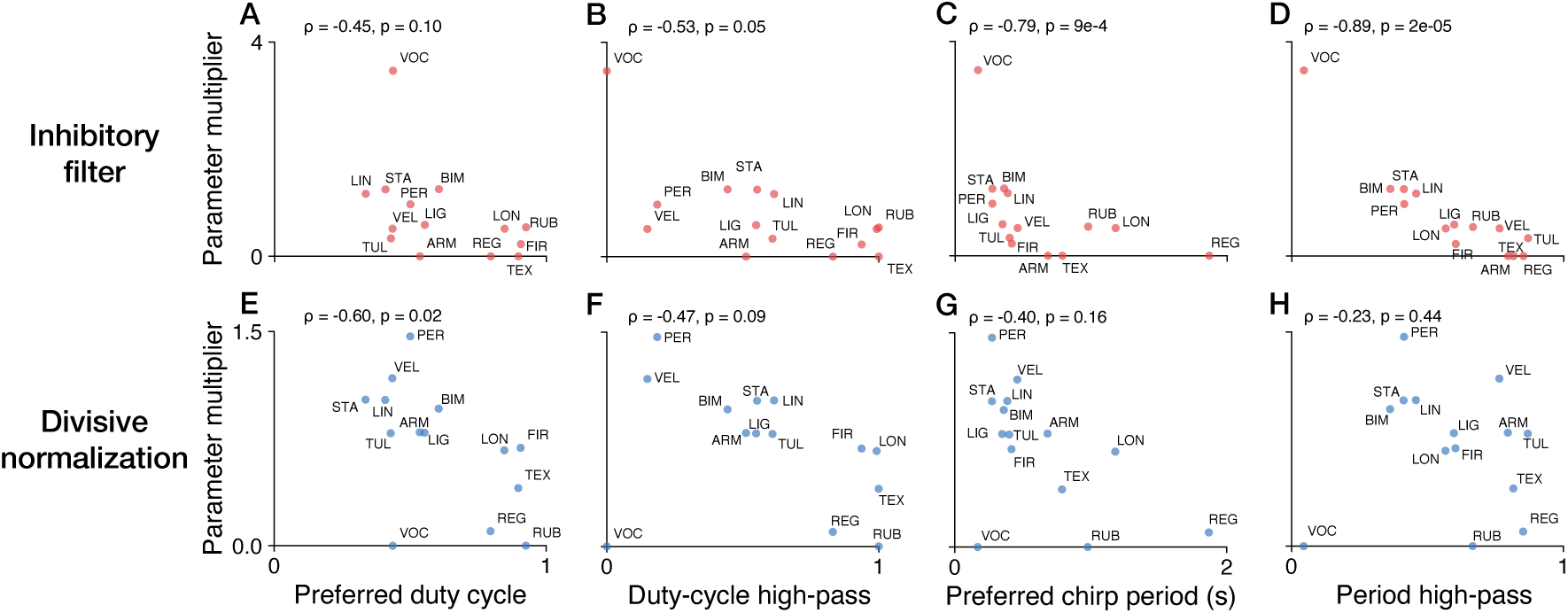
Fitted AN1 parameters and species differences in chirp tuning. A–D. Relationships between the fitted inhibitory-filter multiplier and preferred duty cycle (A), duty-cycle high-pass index (B), preferred chirp period (C), and period high-pass index (D). Each point represents one of the 14 species, labeled with the abbreviation in Table S2. Panel titles give Spearman *ρ* and two-sided *p*-values. **E–H** Same as A–D but for the divisive-normalization multiplier.

